# Natural killer cell function in children with severe malaria differs according to malaria transmission intensity

**DOI:** 10.1101/2025.05.13.653848

**Authors:** Grace Turyasingura, Kagan A. Mellencamp, Ruth Namazzi, Robert O. Opoka, Chandy C. John, Geoffrey T. Hart

**Affiliations:** Department of Microbiology and Immunology, Indiana University School of Medicine, Indianapolis, Indiana, United States; Ryan White Center for Pediatric Infectious Diseases & Global Health, Indiana University School of Medicine, Indianapolis, Indiana, United States; Department of Paediatrics and Child Health, Makerere University, Kampala, Uganda; Division of Infectious Disease and International Medicine, Department of Medicine, University of Minnesota, Minneapolis, Minnesota, United States; Center for Immunology, University of Minnesota, Minneapolis, Minnesota, United States

**Keywords:** *Plasmodium falciparum*, severe malaria, child, natural killer cells, cerebral malaria, severe malarial anemia

## Abstract

Memory-like Natural Killer (NK) cells with enhanced antibody-dependent cellular cytotoxicity (ADCC) have correlated with protection from uncomplicated malaria in prior studies. However, the role of NK cells in severe malaria (SM) has not been characterized. In Ugandan sites with moderate and low malaria transmission, we evaluated NK cell (CD56^bright^, CD56^dim^, CD56^neg^) phenotype and ADCC function by flow cytometry in children <5 years of age with SM (n=21) and control community children (CC, n=19). Children with SM had similar total NK cell counts to CC. Children with SM had a higher proportion of LILRB1+ NK cells than CC. Level of malaria transmission in an area was related to NK cell function. In the low malaria transmission only, children with SM had a higher proportion than CC of NK cells that degranulated, whereas children with SM from both low and moderate malaria transmission areas had lower IFNγ production than CC. We next evaluated functional Boolean gating for degranulation and IFNγ production (CD107a^⁺^/IFNγ^⁻^, CD107a^⁻^/IFNγ^⁺^, and CD107a^⁺^/IFNγ^⁺^) in relation to memory-like and checkpoint/exhaustion NK cell markers in low and moderate malaria transmission SM and CC groups. We found there was a significant increase in degranulating only NK cells (CD107a+, IFNγ−) in children with SM compared to CC solely in the low malaria transmission area. However, there was a significant decrease in NK cells that produced IFNγ but did not degranulate (CD107a−, IFNγ+) in children with SM compared to CC in both low and moderate transmission areas. Our data reveal compound functional differences in NK cells among children with SM living in areas of low versus moderate malaria transmission; however, a consistent finding is reduced NK cell IFNγ production in SM, regardless of transmission intensity.

## INTRODUCTION

*Plasmodium falciparum* malaria is a significant cause of morbidity and mortality, especially in African children under 5 years old(1), who are still developing immunity to the parasite. Severe malaria (SM), marked by vital organ dysfunction, predominantly affects African children. SM can present in many ways, including cerebral malaria, severe malarial anemia, respiratory distress, and acute kidney injury(2–10). Understanding the immune mechanisms that protect against or contribute to SM in children is crucial for effective intervention.

Natural killer (NK) cells, which constitute 5-20% of peripheral blood lymphocytes, influence the immune response through mechanisms such as antibody-dependent cellular cytotoxicity (ADCC) (11–15), natural cytotoxicity (13, 15), and cytokine production(13, 16), especially IFNγ(15–17). NK cells are categorized by CD56 surface expression: CD56^bright^(13), CD56^dim^(13), and CD56^neg^(12, 18). CD56^bright^ NK cells generally have lower cytotoxicity than other subsets, but do effectively produce cytokines(13, 19), while CD56^neg^ NK cells are generally thought to be less functional(18, 20), and CD56^dim^ NK cells typically have the highest cytotoxicity(13, 18, 20). Previous infections, such as cytomegalovirus (CMV), enhance NK cell function, creating adaptive or memory-like NK cells marked by NKG2C and CD57(21, 22), and the loss of the Fc γ chain, PLZF, Syk, and EAT-2(23, 24). Exhaustion and checkpoint inhibitors such as PD-1, CTLA-4, LILRB1, and LAG3 may also affect NK cell function(25–30).

Previous research explored the role of antibody-dependent cellular cytotoxicity (ADCC) in killing infected red blood cells (iRBCs) *in vitro*(*14*). We discovered and now others have replicated that NK cells inhibited iRBC growth via ADCC and released IFNγ(12, 14). Since memory-like NK cells have enhanced ADCC function after CMV infection, we examined memory-like NKG2C+/CD57+ NK cells in Malian children using the additional markers Fc γ chain–, PLZF–, Syk–, and EAT-2–(11, 21–24). As in previous studies, we found no significant increase in the proportion of total NK cells among protected participants(11). This contrasts with other cell subsets and proteins—such as B cells, γδ T cells, CD8 T cells, and antibodies—whose total numbers, memory subsets, or memory-associated proteins (such as IgG), which have been recognized for much longer, have been shown to increase and correlate with protective phenotypes in various malaria contexts(31–46). However, by using recently identified markers of adaptive/memory-like NK cells, we unexpectedly observed a significant increase in the proportion of memory-like CD56^dim^ NK cells among malaria-exposed children and adolescents(11). Among the different memory-like NK cell markers, only Fc γ chain–negative and PLZF–negative NK cells—not NKG2C+/CD57+ NK cells—correlated with resistance to uncomplicated malaria and lower parasitemia(11). Interestingly, these memory-like NK cells show signs of being distinct from NK cells described in non-malaria endemic settings (such as the USA and Europe) that are associated with CMV infection history, as the malaria-exposed memory-like NK cells were predominantly Syk+ and EAT-2+(11, 17). Because we previously showed that antibody-dependent cellular cytotoxicity (ADCC) inhibits the growth of Plasmodium-infected red blood cells in vitro(14), we hypothesize that the enhanced ADCC function of these memory-like NK cells contributes to protection. Increased killing and IFNγ production by these NK cells may increase direct killing of *Plasmodium falciparum* or further activate other immune cells respectively, such as myeloid cells, to help reduce parasitemia.

More recent human studies provide further evidence that NK cells may be protective against uncomplicated malaria. In Ugandan children, high proportions of memory-like CD56^neg^ NK cells were correlated with clinical immunity against uncomplicated malaria, and proportions decreased following reduced malaria exposure(12). In Kenyan children, NK cell ADCC activity against antibody-opsonized *P. falciparum* merozoites correlated with a reduced risk of clinical malaria episodes(47). In mouse models of experimental cerebral malaria, when NK cells are depleted or given no treatment, mice die of experimental cerebral malaria(15). Recent work has shown that when mouse NK cells were stimulated with IL-15 cytokine bound by IL-15Rα/IgG Fc fusion complex, this was sufficient to protect mice from experimental cerebral malaria(46). Although these encouraging human and mouse results suggest a protective role for NK cells, the roles of human CD56^dim^ and CD56^neg^ memory-like NK cells in human severe malaria remain undefined, and the influence of malaria transmission intensity on NK cell function in human severe malaria has not been studied. These gaps highlight the need for focused investigation of human NK cell responses in severe malaria.

Based on the recent human data associating memory-like NK cells markers in CD56^dim^ and CD56^neg^ NK cells and ADCC activity with protection, in the present study we evaluated NK cell phenotype and ADCC function in Ugandan children with severe malaria (SM) and compared them to community children (CC) from the same area with no acute illness. Children were recruited from areas of moderate and low malaria transmission intensity, and differences between SM and CC were compared in both areas. We hypothesized that NK cells in children with SM would exhibit dysfunction, characterized by reduced degranulation, which could lead to less direct killing of iRBCs via ADCC, and reduced IFNγ production, which could decrease stimulation of myeloid cells for phagocytosis or adaptive cells for antibody production. What we found was a complex phenotype where NK functions were both increased and decreased and varied based on disease severity and malaria transmission. These NK functional alterations may lead to an impaired immune response disrupting the balance of clearing *P. falciparum* and maintaining homeostasis, ultimately leading to increased disease severity.

## RESULTS

### Demographic and clinical characteristics of study children

NK cell counts, phenotype and function were evaluated with in children with severe malaria (SM, n=21, 11 with cerebral malaria (CM) and 10 with severe malarial anemia (SMA)) and community children (CC, n=19) of similar age from the same household or neighborhood as the children with SM, with no acute illness or symptoms of disease. Demographic and clinical baseline characteristics of the study children are shown in Table 1. Children with CM had the highest *P. falciparum* parasite density levels, followed by children with SMA and then CC. 9 of the 19 CC had asymptomatic *P. falciparum* parasitemia.

**Table 1.** Baseline demographic and clinical characteristics of study groups.

| Characteristic | CM<br>(N=11) | SMA<br>(N=10) | CC<br>(N=19) | P-value |
| --- | --- | --- | --- | --- |
| Age, years, mean (SD) | 2.2 (0.72) | 1.7 (0.57) | 1.5 (0.56) | **0.01<sup>b</sup>** |
| Sex, male, n (%) | 4 (36.4) | 6 (60.0) | 9 (47.4) | 0.60 |
| Site, Low transmission, n (%) | 6 (54.6) | 3 (30.0) | 12 (63.2) | 0.23 |
| Parasite density, parasites/μL, mean (SD) | 169564 (284399) | 50281 (106474) | 328 (1428) | **<0.001<sup>abc</sup>** |

P-values obtained from an ANOVA for age, Fisher's exact tests for sex and site, and a Kruskal-Wallis test followed by Dunn's test for parasite density. The following groups differed at $p < 0.05$ :
<sup>a</sup> CM vs. SMA; <sup>b</sup> CM vs. CC; <sup>c</sup> SMA vs. CC.
Abbreviations: CM, cerebral malaria; SMA, severe malarial anemia; CC, community children.

### Children with SM had similar absolute NK cell counts to CC, but a higher proportion of LILRB1+ CD56^dim^ and LILRB1+ CD56^bright^ NK cells than CC

When evaluating the number of total NK cells (CD3-, CD7+) and major subsets of NK cells based on CD56 expression, CD56^bright^, CD56^dim^, and CD56^neg^ NK cells, we found no differences in absolute counts of total NK cells or NK cell subsets between children with SM and CC (Figure 1A-E). In addition, we analyzed NK cell subsets in CC with (n=9) vs. without (n=10) asymptomatic *P. falciparum* parasitemia and found no significant differences in total numbers of any cell population or subset (Supplemental Table 1), so we included all CC in the comparative analysis with children with SM.

**Figure 1:**
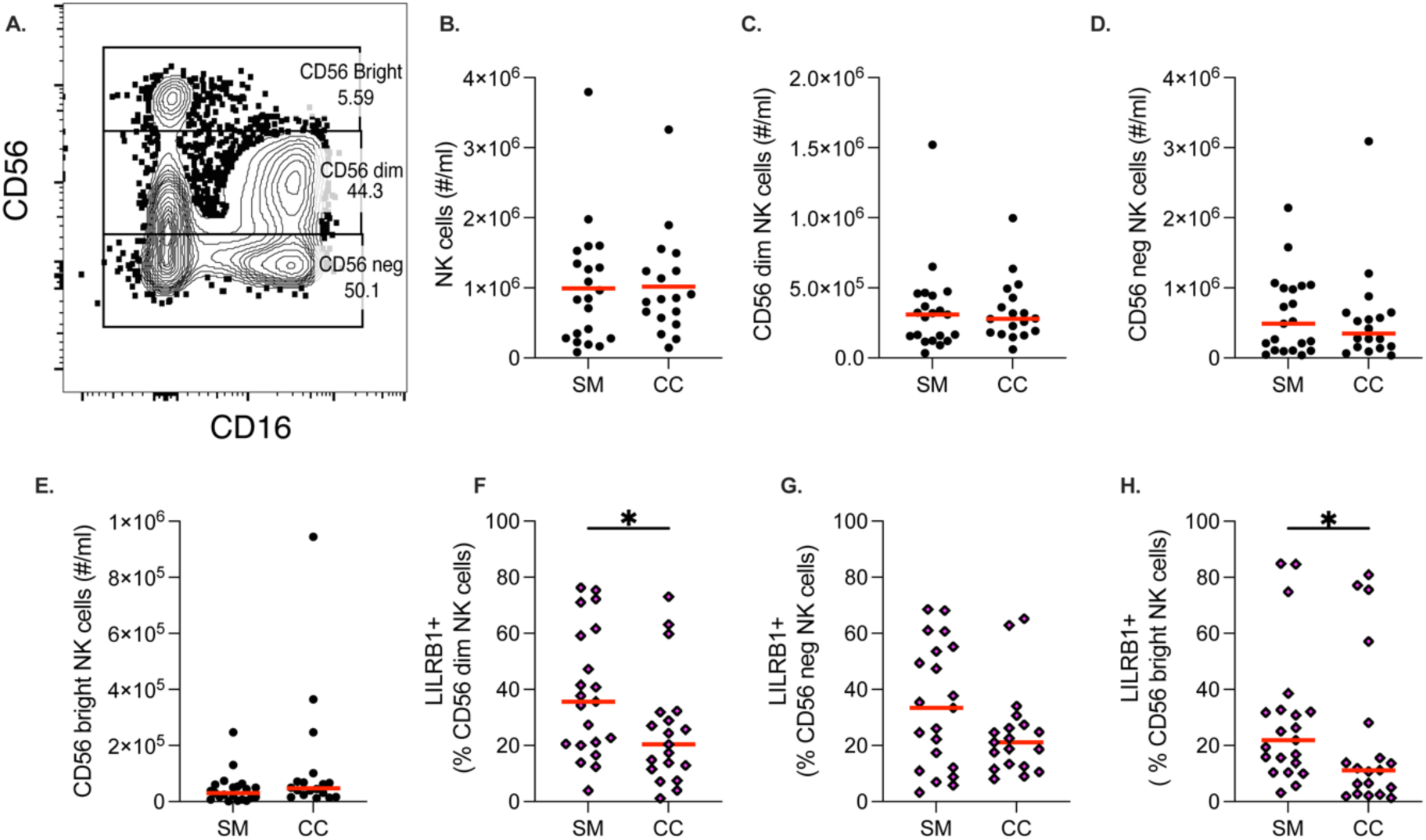
Children with severe malaria (SM) had increased proportion of NK cell inhibitory receptor LILRB1 than community children (CC), but no differences in absolute counts of NK cells. (A) Flow cytometry staining of NK cell subsets defined by Living (Live dead marker and Annexin V(–)), CD64(–), CD3(–), CD7+ showing CD56 and CD16 expression post-ADCC assay. Three major NK cell populations were identified: CD56^bright^, CD56^dim^, and CD56^neg^ after stimulation of PBMCs with uRBCs incubated in anti-human RBC antibody. (B-E) Absolute counts of total NK cells (B). Absolute counts of NK cell subsets: CD56^dim^ (C), CD56^neg^ (D) and CD56^bright^ (E) between SM and CC. (F-G) Analysis of the proportion of LILRB1 in NK subsets: CD56^dim^ (F), CD56^neg^ (G) and CD56^bright^ (H). Each data point represents one participant with the red line indicating the median. Statistical significance was determined using Mann-Whitney U-test analysis between groups SM (n=21), CC (n=18^#^)[^#^One CC was missing the complete blood count (CBC) count] denoted by *p<.05.

To further investigate the phenotypic differences in NK cells, we analyzed NK cell surface markers associated with memory-like NK cells, maturation, activation, and inhibition (e.g., CD57, NKG2C, CD16, LAG3, LILRB1, SIGLEC 7, Fc γ chain (FcRγ)) across four groups: cerebral malaria (CM), severe malarial anemia (SMA), community children (CC), and North American controls (NAC). We found no significant differences in NK cell markers between CM, SMA and CC for CD56^dim^, CD56^neg^, and CD56^bright^ subsets (Supplemental Figure 1A-C). Because we found no major phenotypic differences between the CM and SMA groups, we combined the CM and SMA groups into one severe malaria (SM) group for the remainder of the analysis. When we reanalyzed, the phenotypic data comparing the now combined SM group to CC, we found one marker, LILRB1, was significantly increased in the proportion for both CD56^dim^ and CD56^bright^ NK cell subsets (Figure 1F-H).

### Increased proportion of degranulating NKG2C+, CD57+, LILRB1+, and LAG3+ NK cells in SM

Given that memory-like NK cells demonstrate enhanced ADCC in uncomplicated malaria(11, 12), we evaluated the ADCC function of memory-like NK cell markers in children in the SM and CC groups. To achieve this, we stimulated PBMCs with uninfected RBCs coated with anti-human RBC antibodies and assessed degranulation and IFNγ production within CD56 subsets for the same phenotypic subsets (e.g., CD57, NKG2C, CD16, LAG3, LILRB1, SIGLEC 7, Fc γ chain (FcRγ)). We found that children with SM had a significantly higher proportion of degranulating NKG2C+ NK cells in both CD56^dim^ and CD56^neg^ subsets than CC (Figure 2A-B). Within CD56^dim^ NK cells, the proportion of IFNγ producing NK cells did not significantly differ between the NKG2C populations of SM and CC (Figure 2C). However, among CD56^neg^ NK cells, children with SM had a significantly higher proportion of IFNγ producing NKG2C+ cells than those with CC (Figure 2D).

**Figure 2:**
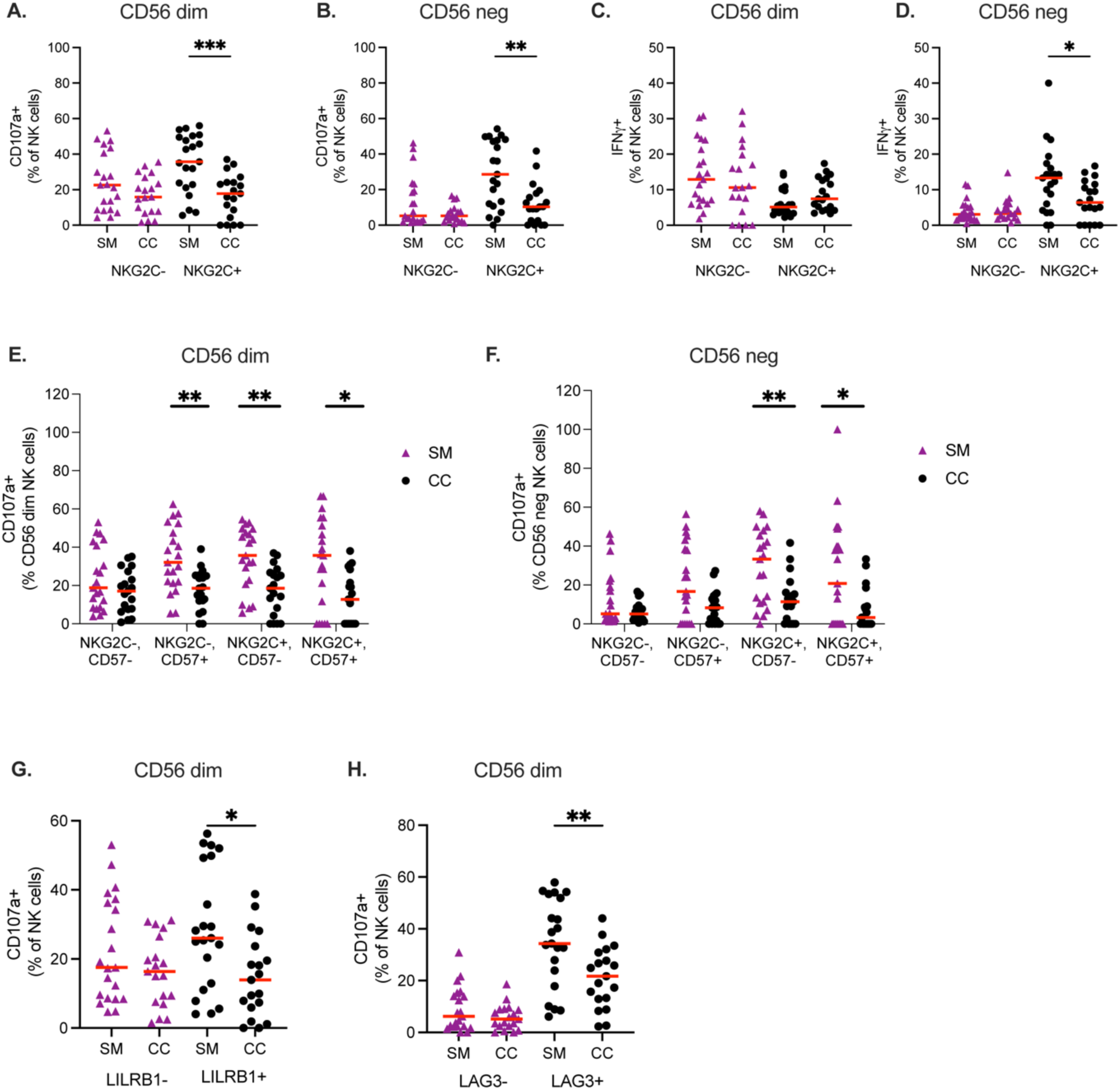
Increased proportion of degranulating NKG2C+, CD57+, LILRB1+, and LAG3+ NK cells in severe malaria (SM) compared to community children (CC) Flow cytometry analysis to evaluate the functional subsets of NK cells after stimulation with uRBCs incubated with anti-human RBC antibody, categorized based on NKG2C and CD57 expression between SM and CC. (A) Proportion of CD107a+ NK cells among NKG2C+ and NKG2C- in the CD56^dim^ NK subset. (B) Proportion of CD107a+ NK cells among NKG2C+ and NKG2C- in the CD56^neg^ NK subset. (C) Proportion of IFNγ producing NK cells among NKG2C+ and NKG2C- in the CD56^dim^ NK subset. (D) Proportion of IFNγ producing NK cells among NKG2C+ and NKG2C- in the CD56^neg^ NK subset. (E) Proportion of CD107a+ NK cells based on NKG2C and CD57 markers in CD56^dim^ NK cells between SM and CC. (F) Proportion of CD107a+ NK cells based on NKG2C and CD57 markers in CD56^neg^ NK cells between SM and CC. (G) Proportion of CD107a+ NK cells in LILRB1+ and LILRB1- populations in CD56^dim^ NK cells. (H) Proportion of CD107a+ NK cells in LAG3+ and LAG3- populations in CD56^dim^ NK cells. Each data point represents one participant with the red line indicating the median. Statistical significance between SM and CC was determined using Mann-Whitney U-test analysis between groups SM (n=21), CC (n=19) denoted by *p<.05, **p<.01, ***p<.001.

Memory-like NK cells, which are increased in CMV-infected individuals, were initially defined as expressing both NKG2C and CD57(21, 22), so we next evaluated the four Boolean populations of NKG2C and CD57 for degranulation and IFNγ production (Figure 2E and 2F). In CD56^dim^ NK cells, the NKG2C−CD57+, NKG2C+CD57−, and NKG2C+CD57+ subsets in children with SM had a higher proportion of degranulation compared to CC (Figure 2E). Similarly, in CD56^neg^ NK cells, NKG2C+CD57− and NKG2C+CD57+ subsets showed an increased proportion of degranulation in SM versus CC (Figure 2F). Lastly, we found that LILRB1+ and LAG3+ CD56^dim^ NK cells had a significantly increased proportion of degranulating (CD107a+) NK cells (Figure 2G and 2H).

### Children with SM in areas of moderate and low transmission demonstrate differences in NK cell function

To assess the impact of malaria transmission on NK cell function in children with SM, we compared NK cell function in children with SM compared to CC in regions of low and moderate malaria transmission. Our analysis revealed there were no significant differences in the proportion of total NK cells in children with SM or CC in moderate or low transmission (Figure 3A). When assessing degranulation function, we found CD56^dim^ but not CD56^neg^ NK cells in the low malaria transmission SM group had higher proportions of degranulating (CD107a+) NK cells relative to both the moderate transmission SM group and low transmission CC group (Figure 3B and C). No difference in IFNγ production between different malaria transmission groups was found for CD56^dim^ or CD56^neg^ NK cells (Figures 3D and E).

**Figure 3:**
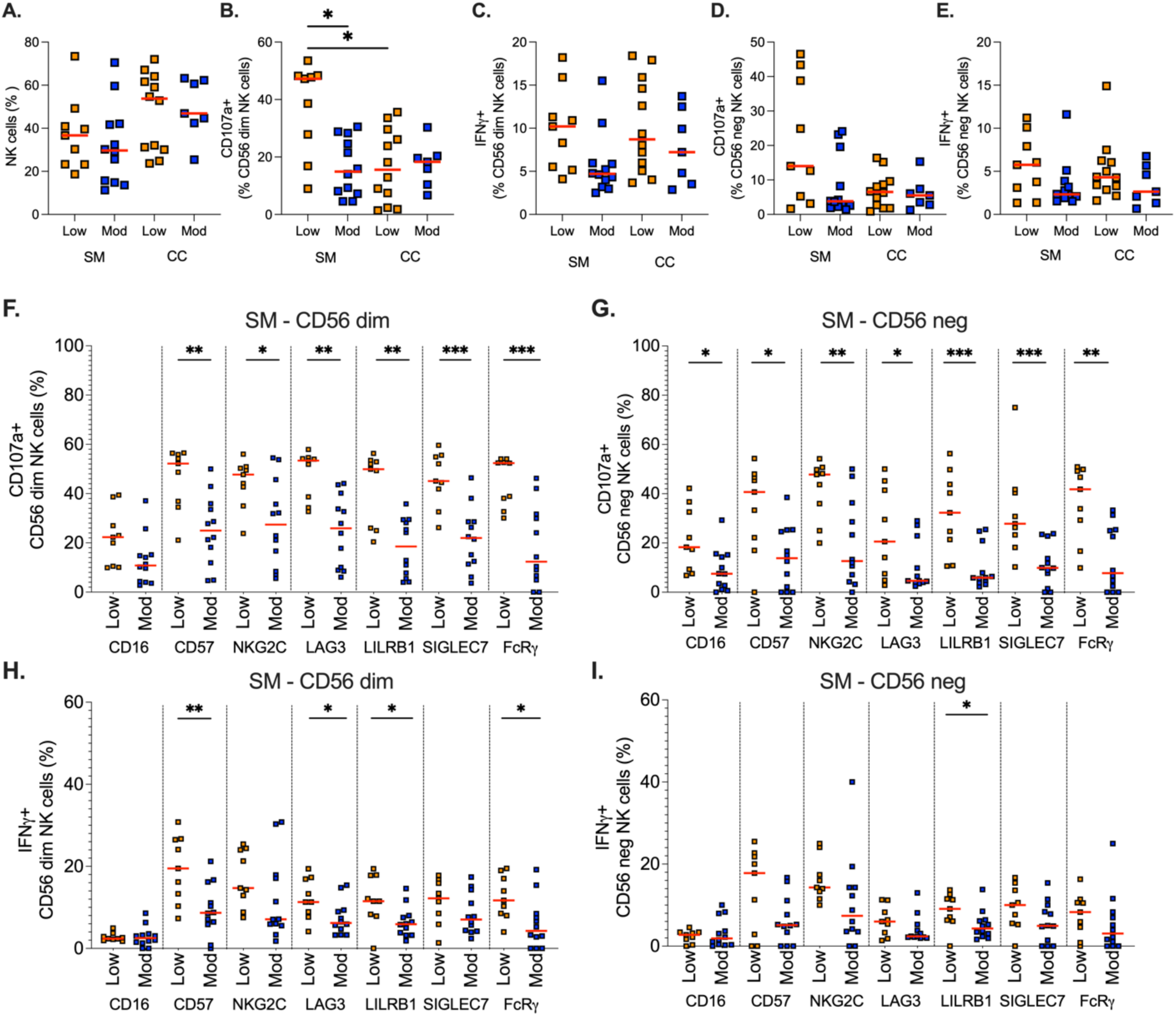
NK cell function in children with severe malaria (SM) differs in areas of low vs. moderate malaria transmission. (A-E) Analysis of NK cell populations for SM and CC in low and moderate malaria transmission sites, when PBMCs were stimulated with uRBCs incubated with anti-RBC antibody. NK cells. (A) Percentage of total NK cells, (B) CD107a+ CD56^dim^ NK cells, and (C) CD107a+ CD56^neg^ NK cells were measured the low and moderate malaria transmission sites. (D-E) Functional analysis of NK cell subsets, focusing on IFNγ production. The percentage of (D) IFNγ+ CD56^dim^ NK cells and (E) IFNγ+ CD56^neg^ NK cells was evaluated under low and moderate malaria transmission sites. (F-G) Analysis of percentage of CD107a in NK cell phenotypic markers for SM (CD16, CD57, NKG2C, LAG3, LILRB1, SIGLEC 7, and FcRγ in CD56^dim^ (F) and CD56^neg^ (G) between low and moderate malaria transmission sites. (H-I) Analysis of percentage of IFNγ in NK cell phenotypic markers for SM (CD16, CD57, NKG2C, LAG3, LILRB1, SIGLEC 7, and FcRγ in CD56^dim^ (H) and CD56^neg^ (I) between low and moderate malaria transmission sites. Each data point represents one participant with the red line indicating the median. Statistical significance between SM and CC in both low and moderate transmission sites (A-E) was determined using pairwise Dunn’s tests following Kruskal-Wallis tests. Statistical significance within SM between low and moderate transmission sites (F-I) was determined using Mann-Whitney U-test analysis. Significance denoted by *p<.05, **p<.01, ***p<.001.

To explore this finding of increased degranulation in the low malaria transmission SM group, we next compared NK function with different NK cell phenotypic markers in children with SM from areas of moderate and low malaria transmission (Figure 3F-G). The analysis revealed a significantly higher proportion of CD107a+ NK cells in both the CD56^dim^ and CD56^neg^ NK subsets for phenotypic markers such as CD57, NKG2C, LAG3, LILRB1, FcRγ, and Siglec 7 in the low malaria transmission area compared to the moderate transmission area (Figures 3F-G). Next, we looked at different phenotypic marker subsets in CD56^dim^ and CD56^neg^ NK cells for IFNγ production (Figure 3H-I). We found a significantly lower proportion of IFNγ+ NK cells for the phenotypic markers CD57, LAG3, LILRB1, and FcRγ in moderate compared to low transmission areas in CD56^dim^ NK cells (Figures 3H), and LILRB1 in CD56^neg^ NK cells. When comparing degranulation and IFNγ production in community children between moderate and low transmission areas, we found no differences in function in different phenotypic subsets (Supplemental Figure 2).

### NKG2C+, CD57+ memory-like NK cells included in cell populations with increased degranulation in low transmission area SM group

We saw an increase proportion of degranulation in low transmission group for NKG2C+ and CD57+ NK cells alone. However, CMV induced memory-like NK cells are defined by NKG2C and CD57 double positive phenotype. Therefore, we assessed the degranulation within low and moderate transmission groups for the Boolean gating scheme of NKG2C and CD57 including NKG2C–/CD57–, NKG2C–/CD57+, NKG2C+/CD57–, and NKG2C+/CD57+ (Figure 4) for both CD56^dim^ (A) and CD56^neg^ (B) NK cells. We found that for CD56dim NK cells, all NKG2C/CD57 Boolean gates in the low transmission group were significantly higher for degranulation in the SM individuals (Figure 4A). For the CD56^neg^ NK cells, all populations that were NKG2C+ had increased degranulation in the low transmission SM group relative to CC (Figure 4B). No significant degranulation was seen with in these Boolean gates in the moderate transmission group between SM and CC individuals (Figure 4A,B).

**Figure 4:**
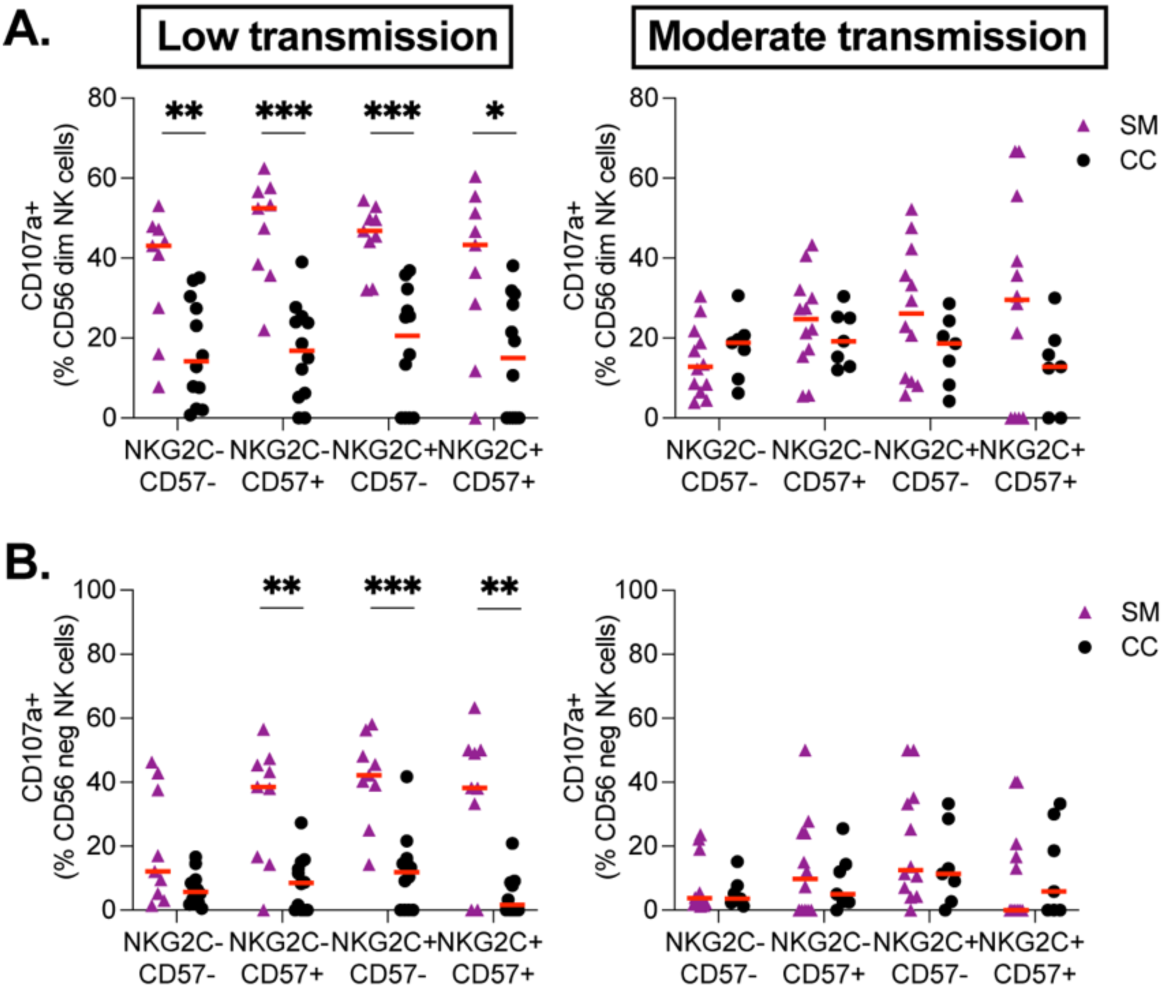
Increased NK cell degranulation in children with severe malaria (SM) compared to community children (CC) is present in low but not moderate malaria transmission areas. In the same generic ADCC assay against RBCs, we separated the individuals by malaria transmission level being low (Left) and moderate (Right). (A) CD56 dim and (B) CD56 neg NK cells for the dual marker Boolean gating of NKG2C and CD57 assessing degranulation (CD107a) after a RBC ADCC assay with an anti-RBC antibody. Each data point represents one participant with the red line indicating the median. Statistical significance between SM and CC was determined using Mann-Whitney U-test analysis between groups SM and CC denoted by *p<.05, **p<.01, ***p<.001.

### Increased proportion of CD107a+IFNγ− and reduced CD107a−IFNγ+ cells in multiple NK cell subsets in children with severe malaria compared to community children

For low and moderate malaria transmission groups, we next evaluated the function of NK cells that would only degranulate, only produce IFNγ, or do both. To do this we performed combinatorial Boolean gating based on CD107a and IFNγ expression, focusing on CD56^dim^ NK cell subsets. This analysis was repeated on PBMCs stimulated with uRBCs incubated with an anti-human RBC antibody. Within either low or moderate transmission groups, three functional groups were identified: degranulation only (CD107a+, IFNγ–), only producing IFNγ (CD107a–, IFNγ+), and both (CD107a+, IFNγ+) (Figure 5). Within the CD56^dim^ degranulation only (CD107a+, IFNγ−) NK cell subset, children in the low transmission group with SM had a significantly higher proportion of degranulation in several markers compared to CC, including NKG2C+, CD57+, LILRB1+, LAG3+, and Siglec 7– (Figure 5A-F (left)). No significant difference was seen in CD56^dim^ degranulating only cells for the moderate transmission group (Figure 5A-E). For the low transmission group within the CD56^dim^ dual functionality group (CD107a+, IFNγ+), there was a significant increase in the NKG2C+ population observed between SM and CC groups (Figure 5A). No other significant differences were found for this dual functioning population for any other marker for low or moderate transmission groups. Finally, within the CD56^dim^ IFNγ only producing group (CD107a−, IFNγ+), children with SM had a lower proportion of IFNγ producing cells in specific markers, including CD57+, LILRB1+, and Siglec 7– for the low transmission group and CD57+ and FcRγ+ for the moderate transmission group (Figure 5A-F) compared to CC.

**Figure 5:**
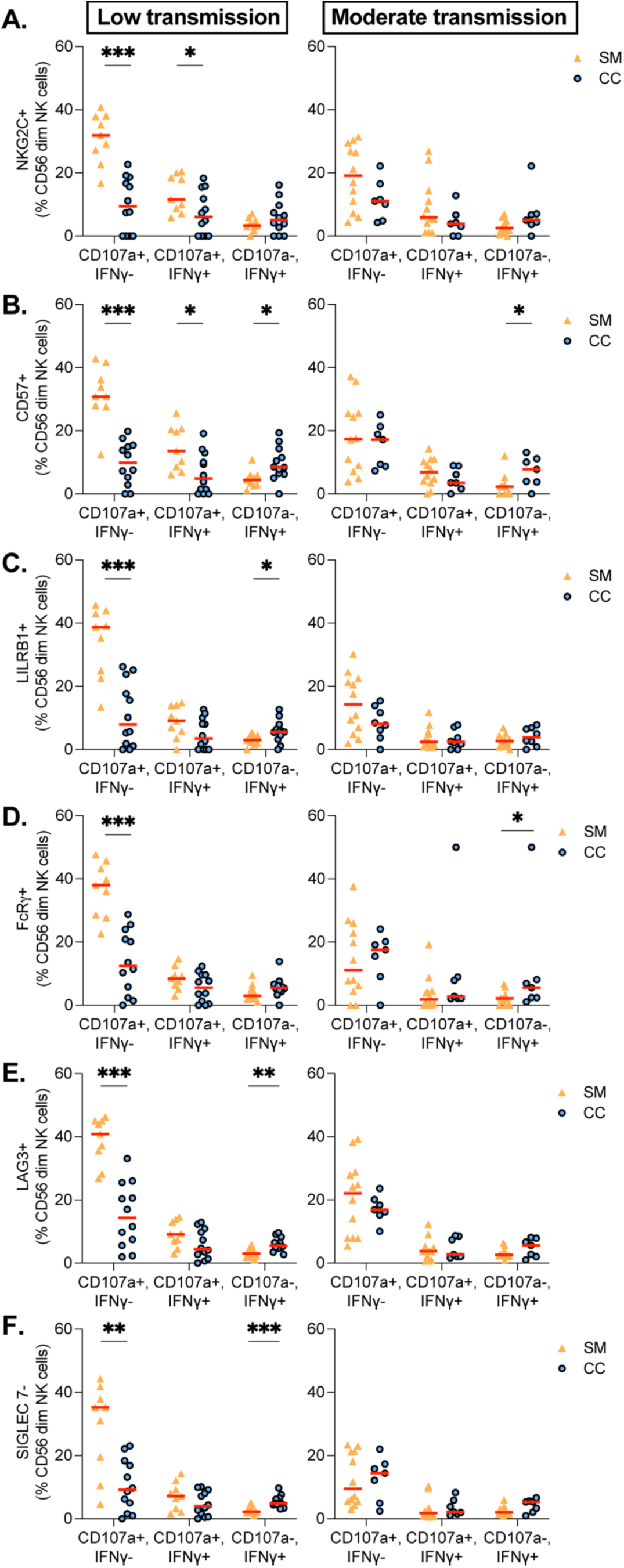
The proportion of CD107a+ IFNγ− NK cell subsets is increased in children with severe malaria (SM) compared to community children (CC) only in an area of low malaria transmission, while the proportion of CD107a− IFNγ+ NK cell subsets is decreased in children with SM compared to CC in low and moderate transmission areas. Data assessing CD107a (degranulation) and IFNγ (cytokine production) when PBMCs were stimulated with uRBCs incubated with anti-RBC antibody. NK cells. CD56^dim^ NK cells were categorized into three populations: CD107a+ IFNγ−, CD107a+ IFNγ+, and CD107a− IFNγ+. (A-H) For low (left) and moderate (right) malaria transmission, percentage of CD107a+ IFNγ−, CD107a+ IFNγ+, and CD107a− IFNγ+ cells in NK cell markers: (A) NKG2C, (B) CD57, (C) LILRB1, (D) FcRγ, (E) LAG3, and (F) SIGLEC 7 neg within the CD56^dim^ NK cell between SM and CC. Each data point represents one participant with the red line indicating the median. Statistical significance between SM and CC was determined using Mann-Whitney U-test analysis between groups SM (n=21), CC (n=19) denoted by *p<.05, **p<.01, ***p<.001.

Overall, the study results reveal that NK cells in children with severe malaria have increased ADCC degranulation and reduced IFNγ production for specific NK cell subsets and this ADCC function also varies according to malaria transmission intensity.

## DISCUSSION

The results of our study offer novel insights into the phenotype and function of NK cells in pediatric severe malaria and highlight how malaria transmission intensity may influence the NK cell immune response in children under five years of age.

Our comparative phenotypic analysis, following PBMC stimulation with opsonized uninfected red blood cells, revealed similar absolute cell numbers of total NK cells and NK cell subsets (CD56^dim^, CD56^bright^, and CD56^neg^). Additionally, our phenotypic analysis showed an increase in the proportion of the inhibitory receptor LILRB1 in the NK cells of children with severe malaria (Figure 1F-1H). LILRB1 is typically thought of as an inhibitory receptor that interacts with HLA class I molecules to suppress NK cell activation, contributing to a hypo- responsive or “exhausted” NK cell phenotype. However, recently, in the context of malaria, LILRB1 has been shown to bind a subset of specific *Plasmodium falciparum-*infected RBC surface proteins called repetitive interspersed repeats (RIFINs). It is hypothesized that when the specific RIFINs that bind to LILRB1 are expressed, this reduces NK cell cytotoxicity and subsequently limits parasite clearance(48, 49). A recent study demonstrated a significant increase in the binding of LILRB1 to infected red blood cells in pediatric patients with severe malaria(48). Our results provide additional evidence to corroborate this finding, where increased LILRB1 on the NK cell surface would theoretically increased binding of infected RBC (iRBC), if the iRBC expresses the corresponding RIFIN proteins. In our study, we investigated LILRB1 using ADCC assays with uninfected RBCs that do not have RIFINs on their surface. It remains unknown whether in children with SM, this increase in LILRB1 would decrease the NK cell functional response when RIFINs on the iRBC surface bind to LILRB1. Interestingly, our results also showed that LILRB1+ NK cells in children with SM had a significantly higher proportion of degranulating NK cells (Figure 2G) in an ADCC assay of uninfected RBCs (RIFIN-negative). This corroborates the results of a separate study showing that an increase in LILRB1+ NK cells was associated with an enhanced ADCC response(12). Further investigation of this complex biology is needed to determine the overall ADCC resulting NK cell function when all these seemingly opposite directional effects are assessed in a controlled way for pediatric severe and uncomplicated malaria.

Building on prior observations associating increased memory-like NK cell functionality with protection in uncomplicated malaria(11, 12), we hypothesized that NK cells in children with severe malaria would lack protective function and exhibit overall dysfunction in both degranulation and IFNγ production compared to community children. Surprisingly, our results revealed significant increases in degranulation and a significant decrease in IFNγ production for different NK cell subsets (Figure 2-5). In addition, when assessing varied malaria transmission groups within the functional data, there were significant NK cell subset increases in degranulation for the SM low transmission group and significant decreases in IFNγ production for NK cell subsets in the SM in both low and moderate malaria transmission group (Figure 3-5).

These functional differences in degranulation were observed in a broad array of memory-like NK cell markers as well as checkpoint molecules, including NKG2C, CD57, LAG3, LILRB1, FcRγ, and Siglec 7 populations. The impact of these functional differences on immune responses in severe malaria requires further investigation. Increased NK cell degranulation could reflect a more robust ADCC response to *Plasmodium* infection; however, excessive degranulation could also indicate excessive cytotoxicity to bystander target cells, causing pathology. Alternatively, because limited exposure to different variants of *P. falciparum* may lead to a lack of sufficient antibody repertoire to infected RBC surface and/or merozoite antigens in children with SM, having the ability to have increased ADCC degranulation without a sufficient antibody repertoire may not be helpful. i.e., increased ADCC is only helpful when the right antibodies are present as well. In this scenario, considering a possible lack of available *P. falciparum* antibody, the increased degranulation may not help direct parasite/iRBC killing but rather worsen the response through a pathogenic bystander killing mechanism.

However, when looking at memory-like NK cell and checkpoint NK cell markers, we also found many markers had decreased IFNγ producing cells in the severe malaria low and moderate transmission group (Figure 3 and 5). We hypothesize that these decreases in IFNγ production would hinder the stimulation of other immune cells, including macrophages and T cells, which play an important role in phagocytosis and helping increase the antibody response, respectively. IFNγ is important for promoting phagocytosis by myeloid cells and facilitating the development of adaptive immunity. This is demonstrated by many studies where malaria-naïve mice lacking IFNγ failed to control early malaria infection, developing high parasitemia and succumbing to disease(50–53). However, IFNγ also promotes the upregulation of T-bet+, a T-box transcription factor, that aids differentiation of naive CD4+ T cells into Th1 cells and atypical B cells, which are thought to generate a less effective antibody responses(31, 32, 54). Therefore, reduced IFNγ production by NK cells may enhance antibody recall responses, but the accompanying decrease in innate phagocytic activity could acutely impair control of parasitemia.

The mechanism of why NK cells have these opposite functional responses with increased degranulation and decreased IFNγ production is unknown. We previously found that CRISPR ablation of Syk increased both degranulation and IFNγ production(17). Ablation of Zap70 decreased both degranulation and IFNγ production. Lastly, ablation of SHP-1 lead to increased degranulation and decreased cytokine production(17). It is unclear if similar proximal CD16 pathway alterations are occurring in these children with severe malaria and vary based on malaria transmission or a more distal mechanisms of differing exhaustion or metabolic differences.

Study limitations included a relatively small sample size, and lack of evaluation of inhibition of red blood cells infected with the strain of parasite infected the child during the severe malaria episode. These limitations are inherent to studies in young children with severe malaria, where volume of sample and the challenge of concurrently culturing the infecting parasite limit the testing that can be done. Study strengths include the first evaluation of NK cell phenotype and function in severe malaria; rigorous study characterization of CM and SMA; the presence of a comparator group of community children to be able to attribute findings to SM; and the evaluation of NK cell function in children with SM and CC in two areas of differing malaria transmission, to determine how differences in transmission may relate to NK cell phenotype and function.

Why NK cell function varies by malaria transmission area is unclear. It is known from our work in uncomplicated malaria that proportion of memory-like NK cells significantly increases with age/exposure(11). Recently it was found that when malaria exposure decreased because of bed net implementation, the proportions of memory-like NK cells populations decreased, particularly memory-like CD56^neg^ FcRγ-negative NK cells(12). These results suggest that malaria infection itself drives the expansion of particular NK cell subsets and can change their function. We hypothesize the cytokine milieu in uncomplicated malaria can stimulate the NK cells alone or in combination with CMV reactivation, that peripherally trains these NK cells to increase in number and have enhanced function broadly. However, in severe malaria this cytokine milieu may be for many reasons different creating a different pathological peripheral training of these NK cells(23, 55). In a recent Covid-19 study on multisystem inflammatory syndrome in children (MIS-C), we found similar to others that peripheral cytokines, such as IL-6, correlate with reduced NK function(56). In this study, both degranulation and cytokine production of NK cells were decreased, supporting the idea that pathogenic strong immune responses correlate with altered NK cell function. Whether this altered NK cell function relates to disease causality could not be established by this study with samples from only the disease time point. Two other major factors that could potentially contribute to site differences in the immune response are co-infection with stool helminths or presence of severe acute malnutrition. These were unlikely to have affected immune responses in the present study because stool helminth infection was rare in study participants (<5% in both sites, manuscript submitted) and severe acute malnutrition was an exclusion criteria for the study. For this reason, differences in malaria transmission are the most likely explanation for the differences in NK cell responses in children in the two study sites.

In conclusion, this study provides novel evidence of altered NK cell function in children with severe malaria, which may be affected by malaria transmission intensity. Our findings highlight key differences in NK cell function in severe malaria in low and moderate transmission. The results raise questions about whether increased NK cell degranulation in low transmission areas contributes to a helpful or pathologic immune response. Similarly, it is not known if the reduced NK cell INFγ production is helpful or pathological in low transmission areas. Further studies evaluating additional immune and parasite parameters over time in children with severe malaria in low and moderate transmission areas are needed to gain further insight into the complexities of NK cell function and its effects in severe malaria. The study findings also highlight the importance of accounting for variations in immune response according to malaria transmission intensity when developing malaria vaccines. Collectively, these results provide a foundation for future studies exploring NK cell research and therapies that may help reduce morbidity and mortality rates associated with severe malaria in children.

## MATERIALS AND METHODS

### Sex as a biological variable

The parent study design enrolled both male and female children, and we conducted specific analysis to investigate if NK cell responses were different between male and females. No differences were found between male and female children in the study.

### Study population

Between 2014 and 2017, we prospectively enrolled 600 children with severe malaria and 120 community children between 6 months to 4 years of age at two sites in Uganda with low and moderate malaria transmission intensity(2, 57). The study was designed to assess cognitive outcomes following severe malaria at 12 months of follow-up in children <5 years of age. Children were recruited at two referral hospitals in central and eastern Uganda: Mulago National Referral Hospital in Kampala and Jinja Regional Referral Hospital in Jinja. Jinja is considered a peri-urban district and has a higher malaria endemicity than Kampala, an urban setting. Children with severe malaria were eligible if they had diagnostic evidence of malaria with either a positive rapid diagnostic test (RDT) for *Plasmodium falciparum* histidine-rich protein 2 (HRP-2) or direct visualization of parasites by Giemsa microscopy as previously described(58). Cerebral malaria was characterized by *P. falciparum* parasitemia and coma, with no other known reason for coma(59). Severe malarial anemia was characterized by *P. falciparum* parasitemia with a hemoglobin level of ≤5 g/dL(60). We focused on CM and SMA because they are the best characterized forms of severe malaria, and are the deadliest and most common form, respectively(61). Exclusion criteria for children with severe malaria included a history of chronic illness requiring medical care (including known HIV infection), history of coma, head trauma, known developmental delay, cerebral palsy, or prior hospitalization for malnutrition.

Community children (CC) were recruited as controls from the extended family or household area of children with severe malaria. Inclusion criteria included residence in the same or nearby neighborhood as a child with severe malaria. Community children were enrolled only if they had no acute illness. Other exclusion criteria for the community children included an illness requiring medical care within the previous 4 weeks, a major medical or neurologic abnormality at screening physical examination, and an active illness or axillary temperature on screening >37.5°C. For the present study, after other sample quality control criteria described below, we evaluated NK function of 19 community children as controls (CC, n=19).

### Study procedures

On enrollment, children had a complete history and physical examination including a blood draw for clinical and study-related procedures. Children with severe malaria were managed according to the Uganda National Guidelines for the treatment of severe malaria, which included intravenous artesunate at a dose of 2.4 mg/kg if ≥20 kg body weight or 3 mg/kg for children <20 kg at 0, 12, and 24 hours after enrollment followed by once-daily intravenous artesunate until the child was able tolerate oral artemether-lumefantrine. Nutritional status was assessed using the height-for-age z score with stunting defined as a height-for-age z score <−2. A complete blood count (CBC) was performed on ethylenediaminetetraacetic acid (EDTA)– anticoagulated blood on admission. Peripheral blood smears were used to quantify parasite density using Giemsa microscopy as previously described(58). HIV counseling and testing was conducted for all study participants according to the national testing algorithm.

### Ethical approval and consent to participate

Initial verbal consent from the parents or legal guardians of study participants was obtained for children fulfilling inclusion criteria, since most participants were critically ill and required emergency stabilization. Written informed consent was obtained once the participant was clinically stabilized. Ethical approval was granted by the Institutional Review Boards at Makerere University School of Medicine, the University of Minnesota, and Indiana University, and regulatory approval granted by the Uganda National Council for Science and Technology.

### PBMC isolation and cryopreservation

PBMCs were isolated from whole blood that was collected in lithium-heparin containing vacutainer tubes (BD) and transferred to the laminar flow hood. For each blood sample, an equal volume of diluted phosphate-buffered saline (PBS) (Sigma) was added in a 15ml conical tube and mixed well. The diluted blood was then layered over 3ml of room temperature histopaque (Sigma) in a conical tube without mixing to keep layers separate. The tubes were centrifuged at 350xg for 30 minutes at room temperature with the brake off. During centrifugation, the PBMCs formed a distinct white blood cell layer (buffy coat) between the plasma and RBC pellet, which was collected and transferred into a new labelled sterile 15ml conical tube. The isolated PBMCs were washed twice by adding sterile PBS to bring total volume to 12ml and centrifuged for 15mins each at 250xg. After washing, the PBMCs were resuspended in 1ml PBS for counting. To determine viability yield of the PBMCs in preparation for cryopreservation cell count and viability were determined using 0.4% trypan blue viability dye (Sigma) to determine viability yield of the PBMCs in preparation for cryopreservation.

After counting, the PBMCs samples were washed by centrifugation at 350xg for 10 mins and the PBMC pellet was then resuspended in cold freezing medium consisting of 90% heat inactivated fetal bovine serum (FBS) (Sigma) and 10% dimethyl sulfoxide (DMSO) at a concentration of 5 million cells/ml. The cell suspension was then aliquoted into labelled cryovials, which were placed in a Mr. Frosty™ freezing container filled with isopropanol (VWR). Right after this, the Mr. Frosty™ freezing container was then transferred to a -80° C freezer to achieve gradual cooling for overnight storage. After 24 hours, the cryovials were transferred to liquid nitrogen for long-term storage. The PBMC samples were then shipped on dry ice by Worldwide Courier Service from Kampala, Uganda to Indianapolis, IN, USA. The samples were then transferred to and stored in a liquid nitrogen to be used for subsequent experiments. North American control (NAC) malaria-naive samples were obtained from Memorial Blood Bank (St. Paul, MN). The PBMCs were processed and frozen as previously described except a different but similar salt gradient, Percoll (GE) was used to isolate the PBMCs. NAC samples were shipped from Minneapolis, MN to Indianapolis, IN on dry ice. These NAC PBMCs were then stored in liquid nitrogen before being used in subsequent experiments.

### PBMC sample selection for analysis

From an initial severe malaria cohort of a total of 720 participants, peripheral blood PBMCs were selected from 55 participants across the respective groups: cerebral malaria (CM=15), severe malaria anemia (SMA=15), and community children (CC=25) based on sample availability. To reduce batch effects, the order of the participants in the severe malaria cohort for sample testing for phenotype and function was randomized. Following analysis, samples exhibiting cell viability below 50% and inadequate antibody staining of internal reference samples were excluded, resulting in a final analysis comprising samples from 40 participants: SM, n=21 (CM, n=11, SMA, n=10) and CC, n=19.

### Antibody-dependent cellular cytotoxicity (ADCC) phenotype and function assays

In a single day, a total of 12 PBMC samples were processed, including three identical NAC PBMC samples that were included for each experiment as phenotype and functional staining quality controls. The PBMC samples were thawed, washed, and resuspended in 2mL of culturing media (RPMI, Serum-free hematopoietic cell medium, with L-Glutamine, gentamicin, and phenol red, 20% heat inactivated human male AB serum (Peak Serum) and supplemented with 20ng/ml recombinant human IL-15 (National Cancer Institute)). The PBMCs were subsequently counted and resuspended in culture media at a concentration of 500,000 cells per 100ul (5 million PBMC/ml).

For the generic ADCC, uRBCs were purified from whole blood by leukocyte reduction filtration (Fenwal Inc.) and resuspended at 50% hematocrit in RPMI-1640 and 25 mM HEPES, L-Glutamine and 50 mg/l Hypoxanthine (K-D Medical). Where cell numbers allowed, we did have a PBMC only control. The uninfected red blood cells and *Plasmodium* infected red blood cells were resuspended in culture media to a concentration of 500,000 cells per 100µL respectively. The uRBC suspension was incubated with rabbit anti-human RBC antibodies (Rockland) for 20 minutes at room temperature. Subsequently, the PBMCs were resuspended in a master mix of anti-human CD107a (or LAMP-1) (Biolegend) at 1:100 (1:200 final), Brefeldin A (Sigma-Aldrich) at 2 µg/ml (1 µg/ml final) concentration and Monensin (Sigma-Aldrich) to 2 µM (1 µM final).

Finally, for each condition, 500,000 PBMCs resuspended in master mix were plated into different wells in a 96 V-bottom plate. The ratio of PBMC to target cells was 1:1, totaling 1 million cells per well in the 96 well plate. For the PBMC only negative control, 100 µL of culturing media was added to the 100ul PBMC resuspended in the master mix cell suspension. The 96 well V-bottom plate (Sarstedt) was then centrifuged for 1 min at 100xg before incubating for 4 hours at 37°C, 5% CO_2_. In every experiment, the same two North American controls were included to be used as both phenotypic and functional staining controls to ensure similarity between experiments.

### Antibody staining and flow cytometry

Following the 4-hour incubation period, cells were washed via centrifugation of the plate at 400xg for 4 minutes. The supernatant was subsequently removed, and the cells were resuspended in a master mix comprising PBS and viability dye (Zombie Aqua, Biolegend) and primary antibody surface stains (Supplemental Table 2). The cells were then incubated at room temperature in the dark for 20 minutes. At the end of the 20-minute incubation period, cells were washed with FACS buffer (PBS, 2% FBS, 2 mM EDTA) and centrifuged at 400xg for 5 minutes, after which the supernatant was discarded. Subsequently, the cells were incubated with Annexin V (Biolegend) resuspended in Annexin V binding buffer at room temperature for 15 minutes. Following the 15-minute incubation, cells were washed with FACS buffer (PBS, 2% FBS, 2 mM EDTA) and centrifuged at 400xg for 5 minutes, and the supernatant was discarded.

For intracellular staining, the cells were washed and incubated in FoxP3 nuclear fixation buffer (Biolegend) at 4°C for 30 minutes as a fixation step, in accordance with the manufacturer’s protocol. Subsequently, the cells were washed with permeabilization buffer, centrifuged at 500xg for 4 minutes, and the supernatant was discarded. The cells were then stained with an intracellular antibody master mix in permeabilization buffer. Following this, the cells were incubated at room temperature in the dark for 30 minutes. After incubation, the cells were washed once with FACS buffer and resuspended in FACS buffer for flow cytometry analysis. Samples were run on an Attune NXT flow cytometer.

### Flow cytometry analysis

Analysis of flow data was conducted utilizing FlowJo 10.3.0 (TreeStar). NK cells were identified by gating on lymphocytes by forward side scatter and side scatter. Then gated on negativity for live/dead marker, Annexin V, and CD64 in the same channel (Supplemental Table 2). Next CD3(–) but CD7+ cells were deemed NK cells. Complete Blood Count data (CBC) data for each subject was used to determine absolute NK cell numbers per ml per participant. For the Boolean analysis of phenotype data, positive gates for seven phenotypic markers (CD16, CD57, NKG2C, SIGLEC 7, LAG3, LILRB1, and FcRγ) were determined, and all single and double combinations of these seven markers were tabulated for each subject. For the analysis of functional data, the markers CD107a and IFNγ were evaluated for single and double combinations within each positive and negative population of the seven markers. Furthermore, the four combinations of NKG2C and CD57 (NKG2C–/CD57–, NKG2C–/CD57+, NKG2C+/CD57–, NKG2C+/CD57+) were examined, as NKG2C and CD57 double-positive cells have been identified as adaptive in previous studies(11). Significant results were discussed in the manuscript.

## Statistical analysis

Data were analyzed using STATA v16.1 (StataCorp) and GraphPad Prism v9.0. Nonparametric analyses were used for assessing differences in NK cell function and phenotype across groups. Mann-Whitney U tests were used to compare two groups, and Kruskal-Wallis tests were used to compare more than two groups, with Dunn’s post-hoc tests for pairwise comparisons. Significance denoted by *p<.05, **p<.01, ***p<.001.

## Data availability

The data for this study are available on request from the corresponding author.

## AUTHOR CONTRIBUTIONS

Conceptualization: GTH, CCJ

Study design: RN, ROO, CCJ, GTH

Designed research experiments: GT, CCJ, GTH

Conducted laboratory testing and acquired data: GT

Analyzed and reviewed data: GT, KAM, CCJ, GTH

Wrote the first draft manuscript: GT, GTH

Review and edited the manuscript: GT, KAM, RN, ROO, CCJ, GTH

## ACKNOWLEDGEMENTS

We extend our appreciation to Mulago Hospital, the children’s ward of Jinja Regional Referral Hospital, and Global Health Uganda for facilitating the research. Additionally, we acknowledge the invaluable contribution of study participants, their caregivers, and the dedicated research teams who provided exemplary clinical care and follow-up to all children involved. This investigation was supported by grants from the NIH Fogarty International Center (D43TW010928), the National Institute of Neurological Disorders and Stroke (R01NS055349), the National Institute of Allergy and Infectious Disease (R01AI146031).

## References

1. Venkatesan, P. 2024. The 2023 WHO World malaria report. Lancet Microbe 5: e214.

2. Namazzi, R., R. Opoka, D. Datta, P. Bangirana, A. Batte, Z. Berrens, M. J. Goings, A. L. Schwaderer, A. L. Conroy, and C. C. John. 2022. Acute Kidney Injury Interacts With Coma, Acidosis, and Impaired Perfusion to Significantly Increase Risk of Death in Children With Severe Malaria. Clin Infect Dis 75: 1511–1519.

3. Birbeck, G. L., M. E. Molyneux, P. W. Kaplan, K. B. Seydel, Y. F. Chimalizeni, K. Kawaza, and T. E. Taylor. 2010. Blantyre Malaria Project Epilepsy Study (BMPES) of neurological outcomes in retinopathy-positive paediatric cerebral malaria survivors: a prospective cohort study. Lancet Neurol 9: 1173–1181.

4. Kremsner, P. G., C. Valim, M. A. Missinou, C. Olola, S. Krishna, S. Issifou, M. Kombila, L. Bwanaisa, S. Mithwani, C. R. Newton, T. Agbenyega, M. Pinder, K. Bojang, D. Wypij, and T. Taylor. 2009. Prognostic value of circulating pigmented cells in African children with malaria. J Infect Dis 199: 142–150.

5. Lewallen, S., R. N. Bronzan, N. A. Beare, S. P. Harding, M. E. Molyneux, and T. E. Taylor. 2008. Using malarial retinopathy to improve the classification of children with cerebral malaria. Trans R Soc Trop Med Hyg 102: 1089–1094.

6. Milner, D. A., Jr., C. Valim, R. A. Carr, P. B. Chandak, N. G. Fosiko, R. Whitten, K. B. Playforth, K. B. Seydel, S. Kamiza, M. E. Molyneux, and T. E. Taylor. 2013. A histological method for quantifying Plasmodium falciparum in the brain in fatal paediatric cerebral malaria. Malar J 12: 191.

7. Milner, D. A., Jr., R. O. Whitten, S. Kamiza, R. Carr, G. Liomba, C. Dzamalala, K. B. Seydel, M. E. Molyneux, and T. E. Taylor. 2014. The systemic pathology of cerebral malaria in African children. Front Cell Infect Microbiol 4: 104.

8. Potchen, M. J., S. D. Kampondeni, K. B. Seydel, G. L. Birbeck, C. A. Hammond, W. G. Bradley, J. K. DeMarco, S. J. Glover, J. O. Ugorji, M. T. Latourette, J. E. Siebert, M. E. Molyneux, and T. E. Taylor. 2012. Acute brain MRI findings in 120 Malawian children with cerebral malaria: new insights into an ancient disease. AJNR Am J Neuroradiol 33: 1740–1746.

9. Taylor, T. E. 2009. Caring for children with cerebral malaria: insights gleaned from 20 years on a research ward in Malawi. Trans R Soc Trop Med Hyg 103 Suppl 1: S6–10.

10. Milner, D. A., Jr. 2018. Malaria Pathogenesis. Cold Spring Harb Perspect Med 8.

11. Hart, G. T., T. M. Tran, J. Theorell, H. Schlums, G. Arora, S. Rajagopalan, A. D. J. Sangala, K. J. Welsh, B. Traore, S. K. Pierce, P. D. Crompton, Y. T. Bryceson, and E. O. Long. 2019. Adaptive NK cells in people exposed to Plasmodium falciparum correlate with protection from malaria. J Exp Med 216: 1280–1290.

12. Ty, M., S. Sun, P. C. Callaway, J. Rek, K. D. Press, K. van der Ploeg, J. Nideffer, Z. Hu, S. Klemm, W. Greenleaf, M. Donato, S. Tukwasibwe, E. Arinaitwe, F. Nankya, K. Musinguzi, D. Andrew, L. de la Parte, D. M. Mori, S. N. Lewis, S. Takahashi, I. Rodriguez-Barraquer, B. Greenhouse, C. Blish, P. J. Utz, P. Khatri, G. Dorsey, M. Kamya, M. Boyle, M. Feeney, I. Ssewanyana, and P. Jagannathan. 2023. Malaria-driven expansion of adaptive-like functional CD56-negative NK cells correlates with clinical immunity to malaria. Sci Transl Med 15: eadd9012.

13. Long, E. O., H. S. Kim, D. Liu, M. E. Peterson, and S. Rajagopalan. 2013. Controlling natural killer cell responses: integration of signals for activation and inhibition. Annu Rev Immunol 31: 227–258.

14. Arora, G., G. T. Hart, J. Manzella-Lapeira, J. Y. Doritchamou, D. L. Narum, L. M. Thomas, J. Brzostowski, S. Rajagopalan, O. K. Doumbo, B. Traore, L. H. Miller, S. K. Pierce, P. E. Duffy, P. D. Crompton, S. A. Desai, and E. O. Long. 2018. NK cells inhibit Plasmodium falciparum growth in red blood cells via antibody-dependent cellular cytotoxicity. Elife 7.

15. Burrack, K. S., G. T. Hart, and S. E. Hamilton. 2019. Contributions of natural killer cells to the immune response against Plasmodium. Malar J 18: 321.

16. Artavanis-Tsakonas, K., and E. M. Riley. 2002. Innate immune response to malaria: rapid induction of IFN-gamma from human NK cells by live Plasmodium falciparum-infected erythrocytes. J Immunol 169: 2956–2963.

17. Dahlvang, J. D., J. K. Dick, J. A. Sangala, P. R. Kennedy, E. J. Pomeroy, K. M. Snyder, J. M. Moushon, C. E. Thefaine, J. Wu, S. E. Hamilton, M. Felices, J. S. Miller, B. Walcheck, B. R. Webber, B. S. Moriarity, and G. T. Hart. 2023. Ablation of SYK Kinase from Expanded Primary Human NK Cells via CRISPR/Cas9 Enhances Cytotoxicity and Cytokine Production. J Immunol 210: 1108–1122.

18. Cao, W. J., X. C. Zhang, L. Y. Wan, Q. Y. Li, X. Y. Mu, A. L. Guo, M. J. Zhou, L. L. Shen, C. Zhang, X. Fan, Y. M. Jiao, R. N. Xu, C. B. Zhou, J. H. Yuan, S. Q. Wang, F. S. Wang, and J. W. Song. 2021. Immune Dysfunctions of CD56(neg) NK Cells Are Associated With HIV-1 Disease Progression. Front Immunol 12: 811091.

19. Poli, A., T. Michel, M. Theresine, E. Andres, F. Hentges, and J. Zimmer. 2009. CD56bright natural killer (NK) cells: an important NK cell subset. Immunology 126: 458–465.

20. Muller-Durovic, B., J. Grahlert, O. P. Devine, A. N. Akbar, and C. Hess. 2019. CD56-negative NK cells with impaired effector function expand in CMV and EBV co-infected healthy donors with age. Aging (Albany NY) 11: 724–740.

21. Foley, B., S. Cooley, M. R. Verneris, M. Pitt, J. Curtsinger, X. Luo, S. Lopez-Verges, L. L. Lanier, D. Weisdorf, and J. S. Miller. 2012. Cytomegalovirus reactivation after allogeneic transplantation promotes a lasting increase in educated NKG2C+ natural killer cells with potent function. Blood 119: 2665–2674.

22. Lopez-Verges, S., J. M. Milush, B. S. Schwartz, M. J. Pando, J. Jarjoura, V. A. York, J. P. Houchins, S. Miller, S. M. Kang, P. J. Norris, D. F. Nixon, and L. L. Lanier. 2011. Expansion of a unique CD57(+)NKG2Chi natural killer cell subset during acute human cytomegalovirus infection. Proc Natl Acad Sci U S A 108: 14725–14732.

23. Schlums, H., F. Cichocki, B. Tesi, J. Theorell, V. Beziat, T. D. Holmes, H. Han, S. C. Chiang, B. Foley, K. Mattsson, S. Larsson, M. Schaffer, K. J. Malmberg, H. G. Ljunggren, J. S. Miller, and Y. T. Bryceson. 2015. Cytomegalovirus infection drives adaptive epigenetic diversification of NK cells with altered signaling and effector function. Immunity 42: 443–456.

24. Zhang, T., J. M. Scott, I. Hwang, and S. Kim. 2013. Cutting edge: antibody-dependent memory-like NK cells distinguished by FcRgamma deficiency. J Immunol 190: 1402–1406.

25. Dick, J. K., J. A. Sangala, V. D. Krishna, A. Khaimraj, L. Hamel, S. M. Erickson, D. Hicks, Y. Soigner, L. E. Covill, A. Johnson, M. J. Ehrhardt, K. Ernste, P. Brodin, R. A. Koup, A. Khaitan, C. Baehr, B. K. Thielen, C. M. Henzler, C. Skipper, J. S. Miller, Y. T. Bryceson, J. Wu, C. C. John, A. Panoskaltsis-Mortari, A. Orioles, M. E. Steiner, M. C. Cheeran, M. Pravetoni, and G. T. Hart. 2024. Antibody-mediated cellular responses are dysregulated in Multisystem Inflammatory Syndrome in Children (MIS-C). bioRxiv.

26. Chen, H., Y. Chen, M. Deng, S. John, X. Gui, A. Kansagra, W. Chen, J. Kim, C. Lewis, G. Wu, J. Xie, L. Zhang, R. Huang, X. Liu, H. Arase, Y. Huang, H. Yu, W. Luo, N. Xia, N. Zhang, Z. An, and C. C. Zhang. 2020. Antagonistic anti-LILRB1 monoclonal antibody regulates antitumor functions of natural killer cells. J Immunother Cancer 8.

27. Beldi-Ferchiou, A., M. Lambert, S. Dogniaux, F. Vely, E. Vivier, D. Olive, S. Dupuy, F. Levasseur, D. Zucman, C. Lebbe, D. Sene, C. Hivroz, and S. Caillat-Zucman. 2016. PD-1 mediates functional exhaustion of activated NK cells in patients with Kaposi sarcoma. Oncotarget 7: 72961–72977.

28. Moebius, J., R. Guha, M. Peterson, K. Abdi, J. Skinner, S. Li, G. Arora, B. Traore, S. Rajagopalan, E. O. Long, and P. D. Crompton. 2020. PD-1 Expression on NK Cells in Malaria-Exposed Individuals Is Associated with Diminished Natural Cytotoxicity and Enhanced Antibody-Dependent Cellular Cytotoxicity. Infect Immun 88.

29. Shapiro, R. M., M. Sheffer, M. A. Booker, M. Y. Tolstorukov, G. C. Birch, M. Sade-Feldman, J. Fang, S. Li, W. Lu, M. Ansuinelli, R. Dulery, M. Tarannum, J. Baginska, N. Dwivedi, A. Kothari, L. Penter, Y. Z. Abdulhamid, I. E. Kaplan, D. Khanhlinh, R. Uppaluri, R. A. Redd, S. Nikiforow, J. Koreth, J. Ritz, C. J. Wu, R. J. Soiffer, G. J. Hanna, and R. Romee. 2025. First-in-human evaluation of memory-like NK cells with an IL-15 super-agonist and CTLA-4 blockade in advanced head and neck cancer. J Hematol Oncol 18: 17.

30. Seiffert, S., A. R. Blaudszun, B. Shibru, J. Korfer, U. Kohl, S. Fricke, U. Sack, and A. Boldt. 2025. Differential Expression of Immune Checkpoints TIM-3, LAG3, TIGIT, and Siglec-7 on Circulating NK Cells: Insights from Healthy Donors compared to Gastric Cancer Patients. Oncol Res Treat: 1–29.

31. Obeng-Adjei, N., S. Portugal, P. Holla, S. Li, H. Sohn, A. Ambegaonkar, J. Skinner, G. Bowyer, O. K. Doumbo, B. Traore, S. K. Pierce, and P. D. Crompton. 2017. Malaria-induced interferon-gamma drives the expansion of Tbethi atypical memory B cells. PLoS Pathog 13: e1006576.

32. Obeng-Adjei, N., S. Portugal, T. M. Tran, T. B. Yazew, J. Skinner, S. Li, A. Jain, P. L. Felgner, O. K. Doumbo, K. Kayentao, A. Ongoiba, B. Traore, and P. D. Crompton. 2015. Circulating Th1-Cell-type Tfh Cells that Exhibit Impaired B Cell Help Are Preferentially Activated during Acute Malaria in Children. Cell Rep 13: 425–439.

33. Portugal, S., J. Moebius, J. Skinner, S. Doumbo, D. Doumtabe, Y. Kone, S. Dia, K. Kanakabandi, D. E. Sturdevant, K. Virtaneva, S. F. Porcella, S. Li, O. K. Doumbo, K. Kayentao, A. Ongoiba, B. Traore, and P. D. Crompton. 2014. Exposure-dependent control of malaria-induced inflammation in children. PLoS Pathog 10: e1004079.

34. Tran, T. M., S. Li, S. Doumbo, D. Doumtabe, C. Y. Huang, S. Dia, A. Bathily, J. Sangala, Y. Kone, A. Traore, M. Niangaly, C. Dara, K. Kayentao, A. Ongoiba, O. K. Doumbo, B. Traore, and P. D. Crompton. 2013. An intensive longitudinal cohort study of Malian children and adults reveals no evidence of acquired immunity to Plasmodium falciparum infection. Clin Infect Dis 57: 40–47.

35. Cohen, S., G. I. Mc, and S. Carrington. 1961. Gamma-globulin and acquired immunity to human malaria. Nature 192: 733–737.

36. Ho, M., H. K. Webster, P. Tongtawe, K. Pattanapanyasat, and W. P. Weidanz. 1990. Increased gamma delta T cells in acute Plasmodium falciparum malaria. Immunol Lett 25: 139–141.

37. Roussilhon, C., M. Agrapart, P. Guglielmi, A. Bensussan, P. Brasseur, and J. J. Ballet. 1994. Human TcR gamma delta+ lymphocyte response on primary exposure to Plasmodium falciparum. Clin Exp Immunol 95: 91–97.

38. Hviid, L., J. A. Kurtzhals, D. Dodoo, O. Rodrigues, A. Ronn, J. O. Commey, F. K. Nkrumah, and T. G. Theander. 1996. The gamma/delta T-cell response to Plasmodium falciparum malaria in a population in which malaria is endemic. Infect Immun 64: 4359–4362.

39. Goodier, M., M. Krause-Jauer, A. Sanni, A. Massougbodji, B. C. Sadeler, G. H. Mitchell, M. Modolell, K. Eichmann, and J. Langhorne. 1993. Gamma delta T cells in the peripheral blood of individuals from an area of holoendemic Plasmodium falciparum transmission. Trans R Soc Trop Med Hyg 87: 692–696.

40. Dorfman, J. R., P. Bejon, F. M. Ndungu, J. Langhorne, M. M. Kortok, B. S. Lowe, T. W. Mwangi, T. N. Williams, and K. Marsh. 2005. B cell memory to 3 Plasmodium falciparum blood-stage antigens in a malaria-endemic area. J Infect Dis 191: 1623–1630.

41. Weiss, G. E., B. Traore, K. Kayentao, A. Ongoiba, S. Doumbo, D. Doumtabe, Y. Kone, S. Dia, A. Guindo, A. Traore, C. Y. Huang, K. Miura, M. Mircetic, S. Li, A. Baughman, D. L. Narum, L. H. Miller, O. K. Doumbo, S. K. Pierce, and P. D. Crompton. 2010. The Plasmodium falciparum-specific human memory B cell compartment expands gradually with repeated malaria infections. PLoS Pathog 6: e1000912.

42. Ndungu, F. M., A. Olotu, J. Mwacharo, M. Nyonda, J. Apfeld, L. K. Mramba, G. W. Fegan, P. Bejon, and K. Marsh. 2012. Memory B cells are a more reliable archive for historical antimalarial responses than plasma antibodies in no-longer exposed children. Proc Natl Acad Sci U S A 109: 8247–8252.

43. Weinbaum, F. I., C. B. Evans, and R. E. Tigelaar. 1976. Immunity to Plasmodium Berghei yoelii in mice. I. The course of infection in T cell and B cell deficient mice. J Immunol 117: 1999–2005.

44. Chen, D. H., R. E. Tigelaar, and F. I. Weinbaum. 1977. Immunity to sporozoite-induced malaria infeciton in mice. I. The effect of immunization of T and B cell-deficient mice. J Immunol 118: 1322–1327.

45. Schofield, L., J. Villaquiran, A. Ferreira, H. Schellekens, R. Nussenzweig, and V. Nussenzweig. 1987. Gamma interferon, CD8+ T cells and antibodies required for immunity to malaria sporozoites. Nature 330: 664–666.

46. Burrack, K. S., M. A. Huggins, E. Taras, P. Dougherty, C. M. Henzler, R. Yang, S. Alter, E. K. Jeng, H. C. Wong, M. Felices, F. Cichocki, J. S. Miller, G. T. Hart, A. J. Johnson, S. C. Jameson, and S. E. Hamilton. 2018. Interleukin-15 Complex Treatment Protects Mice from Cerebral Malaria by Inducing Interleukin-10-Producing Natural Killer Cells. Immunity 48: 760–772 e764.

47. Odera, D. O., J. Tuju, K. Mwai, I. N. Nkumama, K. Furle, T. Chege, R. Kimathi, S. Diehl, F. K. Musasia, M. Rosenkranz, P. Njuguna, M. Hamaluba, M. C. Kapulu, R. Frank, C.-S. S. Team, F. H. A. Osier, A. I. Abdi, P. C. Chi, Z. de Laurent, I. Jao, D. Kamuya, G. Kamuyu, J. Makale, L. Murungi, J. Musyoki, M. Muthui, J. Mwacharo, S. Kariuki, D. Mwanga, J. Mwongeli, F. Ndungu, M. Njue, G. Nyangweso, D. Kimani, J. M. Ngoi, J. Musembi, O. Ngoto, E. Otieno, M. Ooko, J. Shangala, J. Wambua, K. S. Mohammed, D. Omuoyo, M. Mosobo, N. Kibinge, S. Kinyanjui, P. Bejon, B. Lowe, K. Marsh, V. Marsh, Y. Abebe, P. F. Billingsley, B. K. L. Sim, S. L. Hoffman, E. R. James, T. L. Richie, A. Audi, F. Olewe, J. Oloo, J. Ongecha, M. O. Ongas, N. Koskei, P. C. Bull, S. H. Hodgson, C. Kivisi, M. Imwong, S. C. Murphy, B. Ogutu, J. Tarning, M. Winterberg, and T. N. Williams. 2023. Anti-merozoite antibodies induce natural killer cell effector function and are associated with immunity against malaria. Sci Transl Med 15: eabn5993.

48. Saito, F., K. Hirayasu, T. Satoh, C. W. Wang, J. Lusingu, T. Arimori, K. Shida, N. M. Q. Palacpac, S. Itagaki, S. Iwanaga, E. Takashima, T. Tsuboi, M. Kohyama, T. Suenaga, M. Colonna, J. Takagi, T. Lavstsen, T. Horii, and H. Arase. 2017. Immune evasion of Plasmodium falciparum by RIFIN via inhibitory receptors. Nature 552: 101–105.

49. Chen, Y., K. Xu, L. Piccoli, M. Foglierini, J. Tan, W. Jin, J. Gorman, Y. Tsybovsky, B. Zhang, B. Traore, C. Silacci-Fregni, C. Daubenberger, P. D. Crompton, R. Geiger, F. Sallusto, P. D. Kwong, and A. Lanzavecchia. 2021. Structural basis of malaria RIFIN binding by LILRB1-containing antibodies. Nature 592: 639–643.

50. Favre, N., B. Ryffel, G. Bordmann, and W. Rudin. 1997. The course of Plasmodium chabaudi chabaudi infections in interferon-gamma receptor deficient mice. Parasite Immunol 19: 375–383.

51. van der Heyde, H. C., B. Pepper, J. Batchelder, F. Cigel, and W. P. Weidanz. 1997. The time course of selected malarial infections in cytokine-deficient mice. Exp Parasitol 85: 206–213.

52. Amani, V., A. M. Vigario, E. Belnoue, M. Marussig, L. Fonseca, D. Mazier, and L. Renia. 2000. Involvement of IFN-gamma receptor-medicated signaling in pathology and anti-malarial immunity induced by Plasmodium berghei infection. Eur J Immunol 30: 1646–1655.

53. Su, Z., and M. M. Stevenson. 2000. Central role of endogenous gamma interferon in protective immunity against blood-stage Plasmodium chabaudi AS infection. Infect Immun 68: 4399–4406.

54. Zander, R. A., R. Vijay, A. D. Pack, J. J. Guthmiller, A. C. Graham, S. E. Lindner, A. M. Vaughan, S. H. I. Kappe, and N. S. Butler. 2017. Th1-like Plasmodium-Specific Memory CD4(+) T Cells Support Humoral Immunity. Cell Rep 21: 1839–1852.

55. Ochando, J., W. J. M. Mulder, J. C. Madsen, M. G. Netea, and R. Duivenvoorden. 2023. Trained immunity - basic concepts and contributions to immunopathology. Nat Rev Nephrol 19: 23–37.

56. Dick, J. K., J. A. Sangala, V. D. Krishna, A. Khaimraj, L. Hamel, S. M. Erickson, D. Hicks, Y. Soigner, L. E. Covill, A. K. Johnson, M. J. Ehrhardt, K. Ernste, P. Brodin, R. A. Koup, A. Khaitan, C. Baehr, B. K. Thielen, C. M. Henzler, C. Skipper, J. S. Miller, Y. T. Bryceson, J. Wu, C. C. John, A. Panoskaltsis-Mortari, A. Orioles, M. E. Steiner, M. C. J. Cheeran, M. Pravetoni, and G. T. Hart. 2024. NK Cell and Monocyte Dysfunction in Multisystem Inflammatory Syndrome in Children. J Immunol 213: 1452–1466.

57. Sarangam, M. L., R. Namazzi, D. Datta, C. Bond, C. P. B. Vanderpool, R. O. Opoka, C. C. John, and A. L. Conroy. 2022. Intestinal Injury Biomarkers Predict Mortality in Pediatric Severe Malaria. mBio 13: e0132522.

58. Das, D., P. Dahal, M. Dhorda, B. W. Citarella, K. Kennon, K. Stepniewska, I. Felger, F. Chappuis, and P. J. Guerin. 2020. A Systematic Literature Review of Microscopy Methods Reported in Malaria Clinical Trials. Am J Trop Med Hyg 104: 836–841.

59. Idro, R., K. Marsh, C. C. John, and C. R. Newton. 2010. Cerebral malaria: mechanisms of brain injury and strategies for improved neurocognitive outcome. Pediatr Res 68: 267–274.

60. Opoka, R. O., A. L. Conroy, A. Waiswa, R. Wasswa, J. K. Tumwine, C. Karamagi, and C. C. John. 2020. Severe Anemia Is Associated with Systemic Inflammation in Young Children Presenting to a Tertiary Hospital in Uganda. Am J Trop Med Hyg 103: 2574–2580.

61. Mutombo, A. M., O. Mukuku, K. N. Tshibanda, E. K. Swana, E. Mukomena, D. T. Ngwej, O. N. Luboya, J. B. Kakoma, S. O. Wembonyama, J. P. Van Geertruyden, and P. Lutumba. 2018. Severe malaria and death risk factors among children under 5 years at Jason Sendwe Hospital in Democratic Republic of Congo. Pan Afr Med J 29: 184.

