## Supplement for "Natural killer cell function in children with severe malaria differs according to malaria transmission intensity"

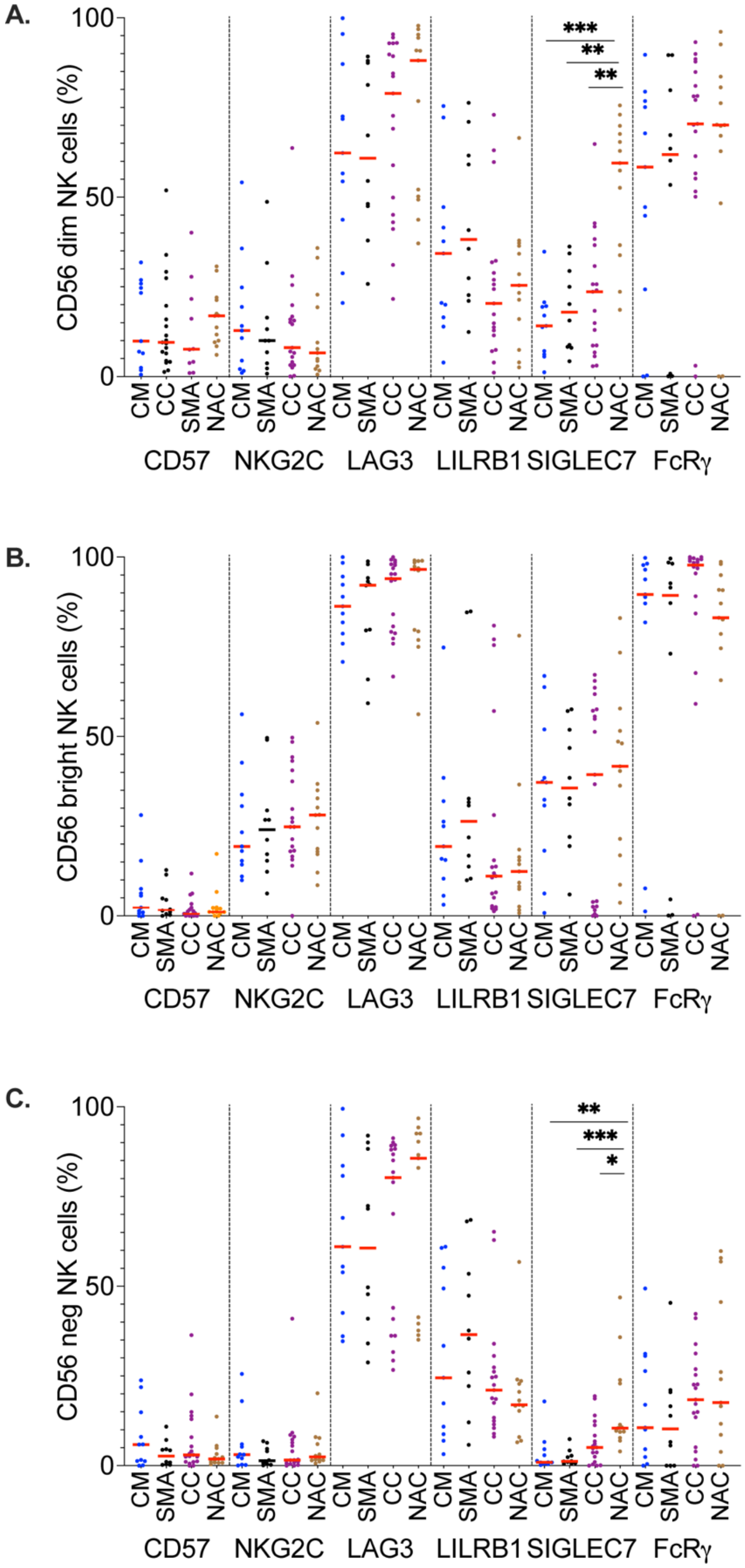
A.
CD56 dim NK cells (%)
***
**
**
B.
CD56 bright NK cells (%)
C.
CD56 neg NK cells (%)
**
***
*
CM
SMA
CC
NAC
CD57
NKG2C
LAG3
LILRB1
SIGLEC7
FcRγ

**Supplemental Figure 1: Phenotypic markers comparison between groups in NK cell subsets** (A-B) Proportions of various NK cell phenotypic markers (CD57, NKG2C, LAG3, LILRB1, SIGLEC 7, and FcRγ) were assesses for each NK subsets when PBMCs were stimulated with uRBCs incubated with anti-RBC antibody: $CD56^{dim}$ (A), $CD56^{bright}$ (B) and $CD56^{neg}$ (C) between different groups (SMA, CM, CC, NAC). Each data point represents one participant with the red line indicating the median. Statistical significance was determined using pairwise Dunn's tests following Kruskal-Wallis tests. Significance denoted by *p<.05, **p<.01, ***p<.001.

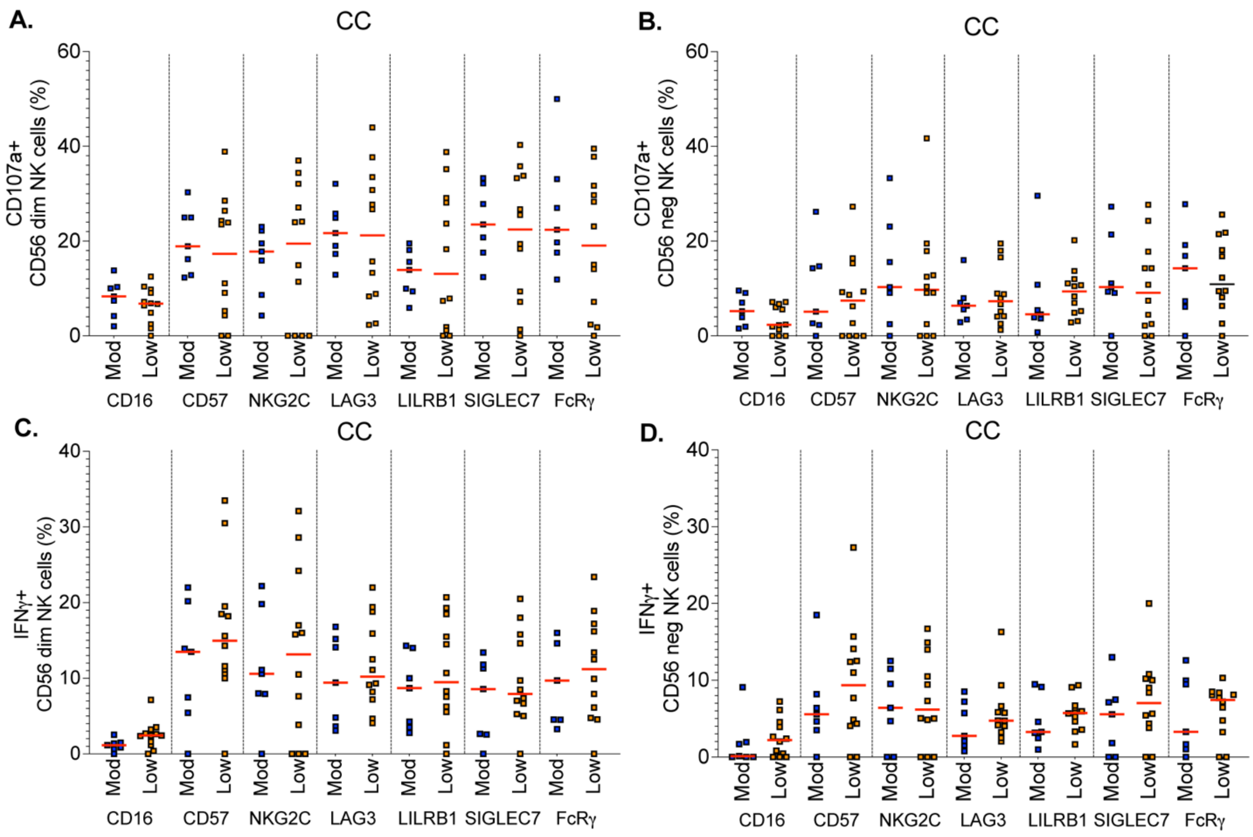


**Supplemental Figure 2: No differences in NK cell function associated with malaria transmission in CC and within site differences.**

(A-D) Analysis of percentage of CD107a (A-B) and IFNγ production (C-D) in NK cell phenotypic markers (CD16, CD57, NKG2C, LAG3, LILRB1, SIGLEC 7, and FcRγ in CD56$^{dim}$ (A,C) and CD56$^{neg}$ (B,D) between low and high malaria exposure sites in community children. Each data point represents one participant with the red line indicating the median. Statistical significance was determined using Mann-Whitney U-test analysis. No significance differences detected.

**Supplemental Table 1.** Absolute counts of NK cells or NK cell subsets in community children with and without asymptomatic *P. falciparum* parasitemia

| **Cell type** | **CC+ (N=9)** | **CC- (N=10)** | **P-value** |
|---|---|---|---|
| NK cells | 6.62 (3.38, 9.10) | 10.47 (6.60, 15.55) | 0.16 |
| $CD56^{dim}$ NK cells | 2.58 (1.80, 3.19) | 2.98 (1.61, 5.24) | 0.55 |
| $CD56^{neg}$ NK cells | 2.71 (0.91, 4.19) | 5.37 (1.66, 6.47) | 0.24 |
| $CD56^{bright}$ NK cells | 0.45 (0.31, 0.71) | 0.44 (0.16, 0.66) | 0.84 |

Data presented as median (IQR) $\times 10^5$/mL. P-values obtained from Mann-Whitney U-tests. Abbreviations: CC+, community children with asymptomatic *P. falciparum* parasitemia; CC-, community children without asymptomatic *P. falciparum* parasitemia.

**Supplement Table 2: Flow cytometry antibodies used**.

| **Antibodies** | **Fluorochrome** | **Clone** | **Vendor** | **Catalog** | **Dilution** |
|---|---|---|---|---|---|
| CD107a | BV421 | H4A3 | Biolegend | 328626 | 1:100 |
| IFNγ | PE-CF594 | B27 | BD | 562392 | 1:50 |
| CD56 | PE-Cy5.5 | A79388 | Beckman | A79388 | 1:50 |
| LAG3 | BV605 | 11C3C65 | Biolegend | 369324 | 1:25 |
| CD16 | BV650 | 3G8 | Biolegend | 302042 | 1:100 |
| CD3 | BV711 | OKT3 | Biolegend | 317328 | 1:100 |
| CD57 | BV780 | QA17A04 | Biolegend | 393328 | 1:100 |
| FcRy | FITC | FCAB5400F | Millipore | FCABS400 | 1:50 |
| CD64 | BV510 | 10.1 | Biolegend | 305028 | 1:100 |
| CD7 | PE | CD7-6B7 | Biolegend | 982706 | 1:100 |
| NKG2C | PE-Cy7 | REA205 | Miltenyi | 130-120-449 | 1:50 |
| LILRB1 | APC | REA998 | Miltenyi | 130-116-616 | 1:50 |
| Siglec 7 | APC-CY7 | REA214 | Miltenyi | 130-101-009 | 1:25 |
| Annexin V | BV510 | | Biologend | 640937 | 1:100 |
| Zombie Aqua | BV510 | | Biolegend | 423102 | 1:500 |
